# Characterizing ketamine-induced dissociation using human intracranial neurophysiology: brain dynamics, network activity, and interactions with propofol

**DOI:** 10.1101/2022.05.02.490333

**Authors:** Fangyun Tian, Laura D. Lewis, David W. Zhou, Gustavo Balanza Villegas, Angelique C. Paulk, Rina Zelmann, Noam Peled, Daniel Soper, Laura A. Santa Cruz Mercado, Robert A. Peterfreund, Linda S. Aglio, Emad N. Eskandar, G Rees Cosgrove, Ziv M. Williams, Robert M. Richardson, Emery N. Brown, Oluwaseun Akeju, Sydney S. Cash, Patrick L. Purdon

## Abstract

Subanesthetic doses of ketamine produce rapid and sustained anti-depressant effects in patients with treatment-resistant depression. Unfortunately, the usefulness of ketamine as a treatment is limited by its potential for abuse because of psychotropic side effects such as dissociation. Understanding the brain dynamics and the neural circuits involved in ketamine’s effects could lend insight into improved therapies for depression with fewer adverse effects. It is believed that ketamine acts via NMDA receptor and hyperpolarization-activated cyclic nucleotide-gated 1 (HCN1) channels to produce changes in oscillatory brain dynamics. Here we show, in humans, a detailed description of the principal oscillatory changes in cortical and subcortical structures by administration of a subanesthetic dose of ketamine. Using recordings from intracranial electrodes, we found that ketamine increased gamma oscillations within prefrontal cortical areas and the hippocampus--structures previously implicated in ketamine’s antidepressant effects. Furthermore, our studies provide direct evidence of a ketamine-induced 3 Hz oscillation in posteromedial cortex that has been proposed as a mechanism for its dissociative effects. By analyzing changes in neural oscillations after the addition of propofol, whose GABAergic activity antagonizes ketamine’s NMDA-mediated disinhibition alongside a shared HCN1 inhibitory effect, we identified brain dynamics that could be attributed to NMDA-mediated disinhibition versus HCN1 inhibition. Overall, our results imply that ketamine engages different neural circuits in distinct frequency-dependent patterns of activity to produce its antidepressant and dissociative sensory effects. These insights may help guide the development of novel brain dynamic biomarkers and therapeutics for depression.

## Main text

Ketamine is a dissociative anesthetic that has both anesthetic and psychoactive properties^1,2^. Intravenous induction doses (1-2mg/kg) of ketamine result in a rapid loss of consciousness appropriate for general anesthesia^3,4^. At subanesthetic doses (0.5 mg/kg), ketamine produces a dissociative state, which includes gaps in memory, out of body experiences, and altered sensory perception^5,6^. In addition, intravenous administration of a subanesthetic dose of ketamine induces significant and rapid anti-depressant-like response in depressed patients^7^. Although ketamine was approved by the Food and Drug Administration (FDA) for adult patients with treatment-resistant depression^8^, the neuropsychiatric side effects have limited its extensive use in clinical practice. Defining the neural circuits engaged in ketamine’s rapid anti-depressant and dissociative effects is an important priority that could facilitate development of improved therapies with fewer side effects and greater safety.

Ketamine is known to induce profound changes in brain oscillatory dynamics that appear to be correlated with its anti-depressant and sensory dissociative activity^9-13^. The electrophysiologic profile of subanesthetic ketamine in humans generally includes an increase of gamma oscillation power and a decrease of delta, alpha, and beta oscillation power^9-12^. Ketamine’s effects on theta power are reported less consistently in the literature^9-12^. Oscillatory power changes have also been reported in patients with depression and have been used to differentiate depressive from healthy subjects^13^. However, the relationships between these changes in oscillatory power and the neural circuit mechanisms of depression and dissociation are not well-understood. Previous studies suggest that at subanesthetic doses, ketamine preferentially blocks the NMDA receptors on GABAergic inhibitory interneurons, resulting in disinhibition of downstream excitatory pyramidal neurons, that is thought to facilitate increased gamma-band activity^14-16^. When GABA_A_ agonists, such as benzodiazepines, are administered alongside ketamine, they mitigate dissociations, possibly by restoring inhibitory activity in the affected brain regions^17,18^. In addition, ketamine inhibits the hyperpolarization-activated cyclic nucleotide-gated potassium channel 1 (HCN1), a molecular target that is thought to play an important role in generating rhythmic EEG activity and is considered a novel therapeutic target for depressive disorders^19-21^. It is, however, unclear which cortical or subcortical structures play a major role in mediating this process. The dissociative state is likely the result of disrupted spatial and temporal coordination between multiple brain regions that normally work in concert during sensory and cognitive processing^18^. Previous work has showed that ketamine’s anti-depressant effects are largely dependent upon its actions within the prefrontal cortex and the hippocampus^22^. On the other hand, the reduction of alpha oscillations in the precuneus and temporal-parietal junction and the 3 Hz rhythm in the deep posteromedial cortex (PMC), as studied in rodents, have been proposed as mechanisms for ketamine-induced dissociation^10,23^. Although ketamine’s anti-depressive and dissociative effects are known to co-occur whenever the drug is administered, these effects may in fact be mediated by distinct mechanisms within distinct neural circuits. If that were true, it might be possible to design novel therapeutics with greater specificity and fewer side effects.

In this study, we measured intracranial EEG (iEEG) in human patients implanted with intracranial electrodes who were administered a subanesthetic dose of ketamine prior to induction of general anesthesia with propofol for electrode removal surgery. To characterize the potential roles of NMDA receptors and HCN1 channels, we also analyzed how propofol, a positive GABA allosteric modulator and HCN1 blocker, changed the ketamine-induced activity. Compared with surface EEG, the iEEG provides greater spatial specificity, which allows us to accurately characterize the dynamics that occur in a variety of cortical and subcortical structures simultaneously during the dissociative state. By comparing the iEEG power changes before and after the administration of ketamine and propofol, we identified brain dynamics that are potentially mediated by NMDA receptor disinhibition distinct from HCN1 inhibition mechanisms. To characterize the role of 3 Hz rhythms in ketamine-induced dissociative state, we also analyzed the spatial distribution of 3 Hz rhythms after ketamine and propofol administration, neither of which to our knowledge has been directly studied in humans previously.

We collected data from 10 epilepsy patients implanted with intracranial depth electrodes to identify sites of epileptogenic origin (**Extended Data Table 1**). The responses on the Clinician-Administered Dissociative States Scale (CADSS) questionnaire are summarized in **Extended Data Table 2**. Among the total of 23 questions, nine out of the ten subjects answered yes to more than 20% of the questions, indicating that they had entered into a dissociative state. The percentage of yes (8.7%-91.3%) and no (0%-91.3%) to questions varies between subjects, suggesting that infusion of a subanesthetic dose (0.5mg/kg over 14 minutes) of ketamine induced a variety of dissociative experiences for different subjects. The responses on the questionnaire confirmed that our subanesthetic ketamine administration paradigm induced a dissociative state.

We observed distinct dynamic patterns in the iEEG after ketamine infusion, which changed after administration of propofol. **Figure 1** shows the spectrogram and power spectra for 3 representative channels in the inferior frontal, middle temporal, and occipital cortices. Under ketamine, we observed increased gamma power (25-55Hz) in the inferior frontal channel and decreased alpha power (8-15Hz) in the middle temporal and occipital channels. After propofol was added, there was a large increase of power in the inferior frontal and middle temporal channels for nearly all frequencies, except for high gamma band (40-55Hz). In contrast, the reduction of alpha oscillations in the occipital channel was further enhanced with the addition of propofol. To understand how these brain dynamics mapped to different brain structures, we analyzed the changes in power for different cortical and subcortical structures, first after ketamine infusion (**Figure 2**, see **Supplementary Figure 2** and **Table 3** for statistics) and then after the addition of propofol (**Figure 3**, see **Supplementary Figure 3** and **Table 4** for statistics).

**Figure 1.**
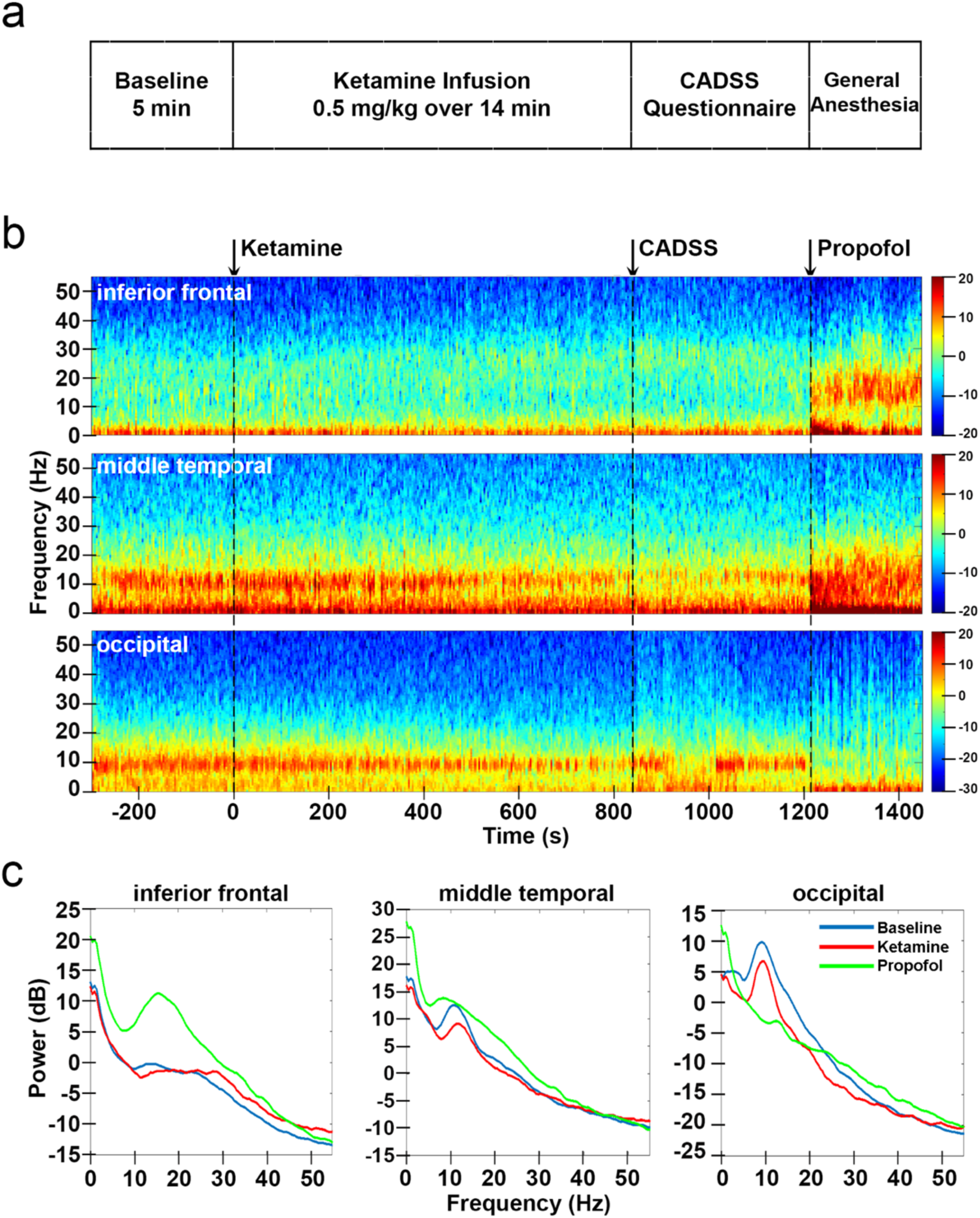
Study protocol and intracranial EEG power changes for representative channels. **a**, study protocol. **b**, power spectrogram (dB) for three representative channels: inferior frontal, middle temporal and occipital. **c**, power spectrum averaged across time during baseline, ketamine, and propofol conditions. CADSS: clinician-administered dissociative states scale.

**Figure 2.**
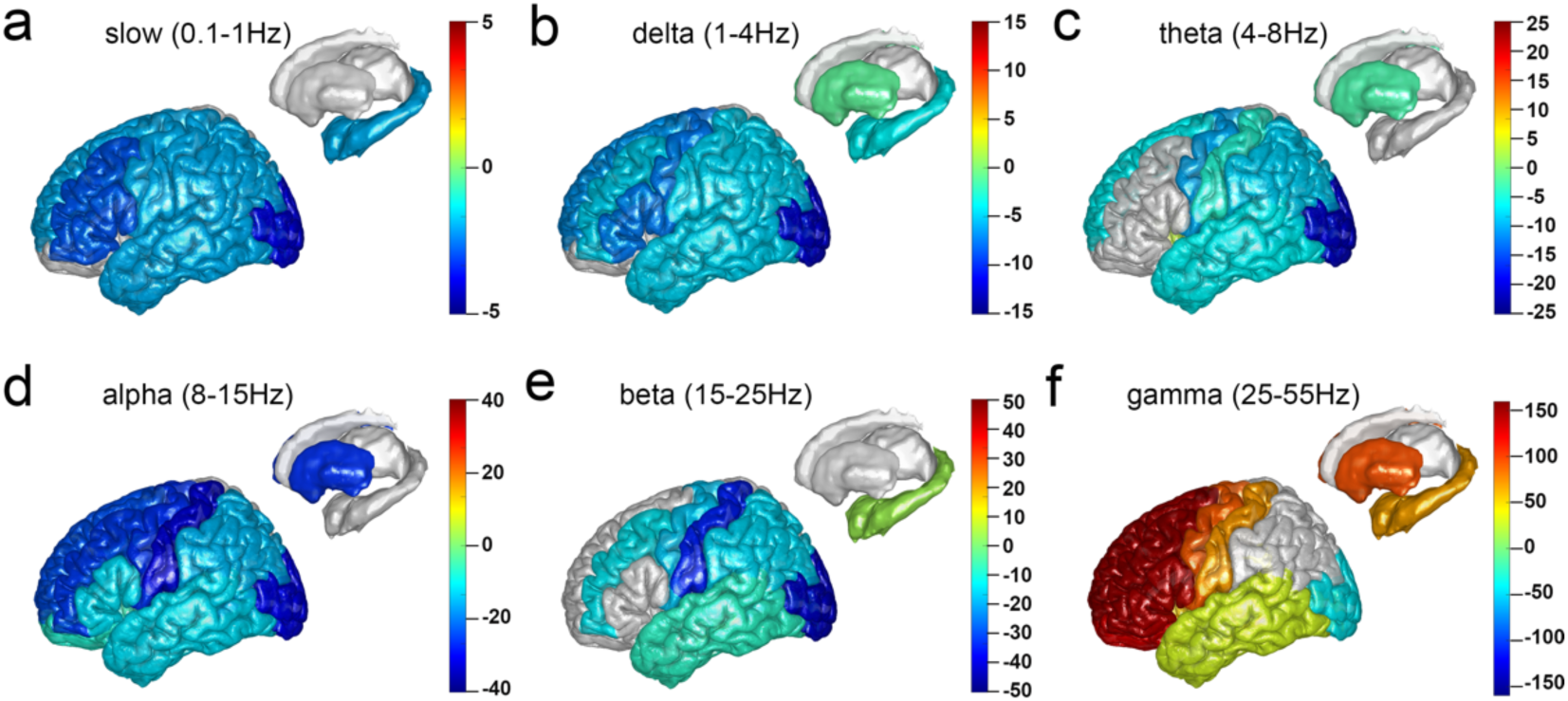
Structural mapping for intracranial EEG power changes after ketamine infusion relative to baseline at 6 frequencies. The mean differences in power (dB) after ketamine infusion relative to baseline were calculated for 824 channels across 10 subjects and plotted on Colin 27 brain template. **a**, slow: 0.1-1Hz. **b**, delta: 1-4Hz. **c**, theta: 4-8Hz. **d**, alpha: 8-15Hz. **e**, beta: 15-25Hz. **f**, gamma: 25-55Hz. Warmer colors indicate a significant increase in power, cooler colors indicate a significant decrease in power, and a grey color indicates no significant change in power (*P* < 0.05).

**Figure 3.**
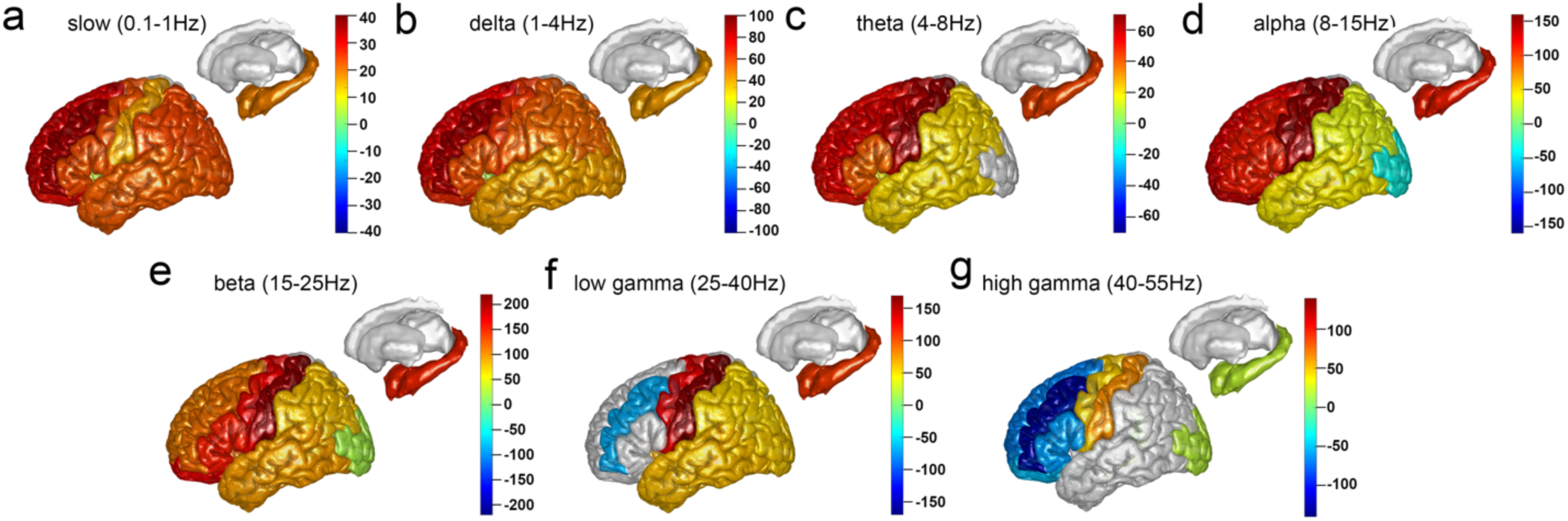
Structural mapping for intracranial EEG power changes after propofol bolus relative to ketamine period at 7 frequencies. The mean differences in power (dB) after propofol bolus relative to ketamine period were calculated for 606 channels across 7 subjects and plotted on Colin27 brain template. **a**, slow: 0.1-1Hz. **b**, delta: 1-4Hz. **c**, theta: 4-8Hz. **d**, alpha: 8-15Hz. **e**, beta: 15-25Hz. **f**, low gamma: 25-40Hz. **g**, high gamma: 40-55Hz. Warmer colors indicate a significant increase in power, cooler colors indicate a significant decrease in power, and a grey color indicates no significant change in power (*P* < 0.05).

For gamma frequencies (25-55 Hz), we observed a remarkable increase in power after ketamine infusion in frontal structures, including the anterior and posterior cingulate, superior frontal, middle frontal, orbitofrontal, and inferior frontal cortex (**Figure 2f**). We also identified mild increases in gamma power in the isthmus cingulate, parietal, temporal, and sensorimotor cortices, the insula, and limbic structures. When propofol was administered (**Figure 3f&g**), it reversed the gamma power increase observed in the frontal cortex. The large increase of gamma power in frontal regions of the brain has been reported with both subanesthetic and anesthetic doses of ketamine and is highly consistent within the literature^9-12,24^. We propose that the ketamine-induced gamma power increase and propofol-induced gamma power reduction in prefrontal regions of the brain may be explained by an antagonist mechanism (**Figure 5, top panel**). Ketamine preferentially blocks the NMDA receptors on GABAergic inhibitory interneurons, resulting in disinhibition of the downstream excitatory pyramidal neurons, which mediates the increased gamma-band activity^14-16^. When propofol, a GABA agonist, is administered alongside ketamine, it antagonizes the gamma power increase by restoring some of the inhibitory activity in the prefrontal cortex. The increase in gamma spectral power anteriorly following subanesthetic ketamine infusion may reflect a shift of brain activity from a globally balanced state to a disorganized and autonomous state^25^. The changes in gamma band activity in sensory cortices may contribute to the discoordination of higher-order functional networks and perceptual distortions produced by subanesthetic doses of ketamine^26-28^.

In contrast, for alpha frequencies (8-15 Hz), we detected a large reduction in iEEG power after ketamine infusion, for all brain regions studied (**Figure 2d**). The largest reductions were observed in postcentral, lingual, pericalcarine, and occipital cortices. The addition of propofol caused a further reduction of alpha power in lingual, pericalcarine, and occipital cortices, whereas the alpha power reduction at all other brain regions was dramatically reversed by propofol (**Figure 3d**). The attenuated alpha power in occipital regions with the addition of ketamine and propofol may be attributed to their shared inhibition at HCN1 channels (**Figure 5, middle panel**). HCN1 channels were identified as an important molecular target for ketamine’s action^19^. Knockout of HCN1 channels abolishes the ketamine-induced loss-of-right reflex, a behavioral correlate of unconsciousness in rodents^19^. Propofol also inhibits HCN1 channels and the HCN1 knock-out mice were found to be less sensitive to unconsciousness due to propofol^19^. As the shared molecular target for ketamine and propofol, HCN1 might account for the additive effects on iEEG power changes when both drugs were administered together. This is also supported by modeling studies which showed that the reduction in hyperpolarization-activated cationic current (*Ih*) mediated by HCN1 abolished occipital alpha rhythm by silencing thalamocortical cells, whereas the increase in GABAergic inhibition modulated the increase of frontal alpha activity^29^. The reduction of alpha power in occipital regions is also observed during anesthetic doses of ketamine^9,11^, propofol-induced unconsciousness^30^, as well as sleep^31,32^, suggesting the loss of occipital alpha rhythms may be a hallmark for disrupted sensory processing^33^.

We studied the spatial distribution of 3 Hz rhythms after the administration of ketamine and propofol (**Figure 4**). We identified a significant increase of 3-4 Hz oscillatory power after ketamine infusion in posterior and isthmus cingulate cortex, which are part of the PMC, as well as the pars opercularis located within the inferior frontal cortex(**Figure 4a&b**). We then analyzed the spectrum of the oscillatory activity within PMC by plotting the power differences after ketamine relative to baseline for posterior and isthmus cingulate cortex as a function of the frequency (**Figure 4c**). We found that a significant increase of iEEG power after ketamine occurred between 3 to 6 Hz, showing a clear oscillatory peak within this frequency range. The addition of propofol significantly increased the 3-4 Hz power in most brain regions, including the posterior and isthmus cingulate cortex, suggesting that the effects of ketamine and propofol on this 3-4Hz rhythm are additive rather than antagonistic (**Figure 4d&e**). Vesuna, et al., 2020, showed that there are NMDA receptors and HCN1 channels in the homologous deep retrosplenial (RSP) cortex in mice, both of which are required for generating the observed 3 Hz rhythmic activity^23^. Knockout of HCN1 channels abolished ketamine-induced rhythm in RSP and the dissociation-related behavior in mice, whereas optogenetic inhibition of long-range inputs to the RSP enhanced ketamine-induced oscillations^23^. Vesuna, et al., proposed that ketamine blockade of NMDA receptors could hyperpolarize membrane potentials in PMC, activating intrinsic HCN1 channels and permitting rhythmic dynamics. We propose that the same effect could occur with propofol by way of a GABA-mediated hyperpolarization (**Figure 5, bottom panel**). Although both ketamine and propofol induced 3Hz rhythms in PMC, dissociation was only detected after ketamine. This may be because propofol suppresses arousal and induces unconsciousness, which would supersede any perceived dissociative effects.

**Figure 4.**
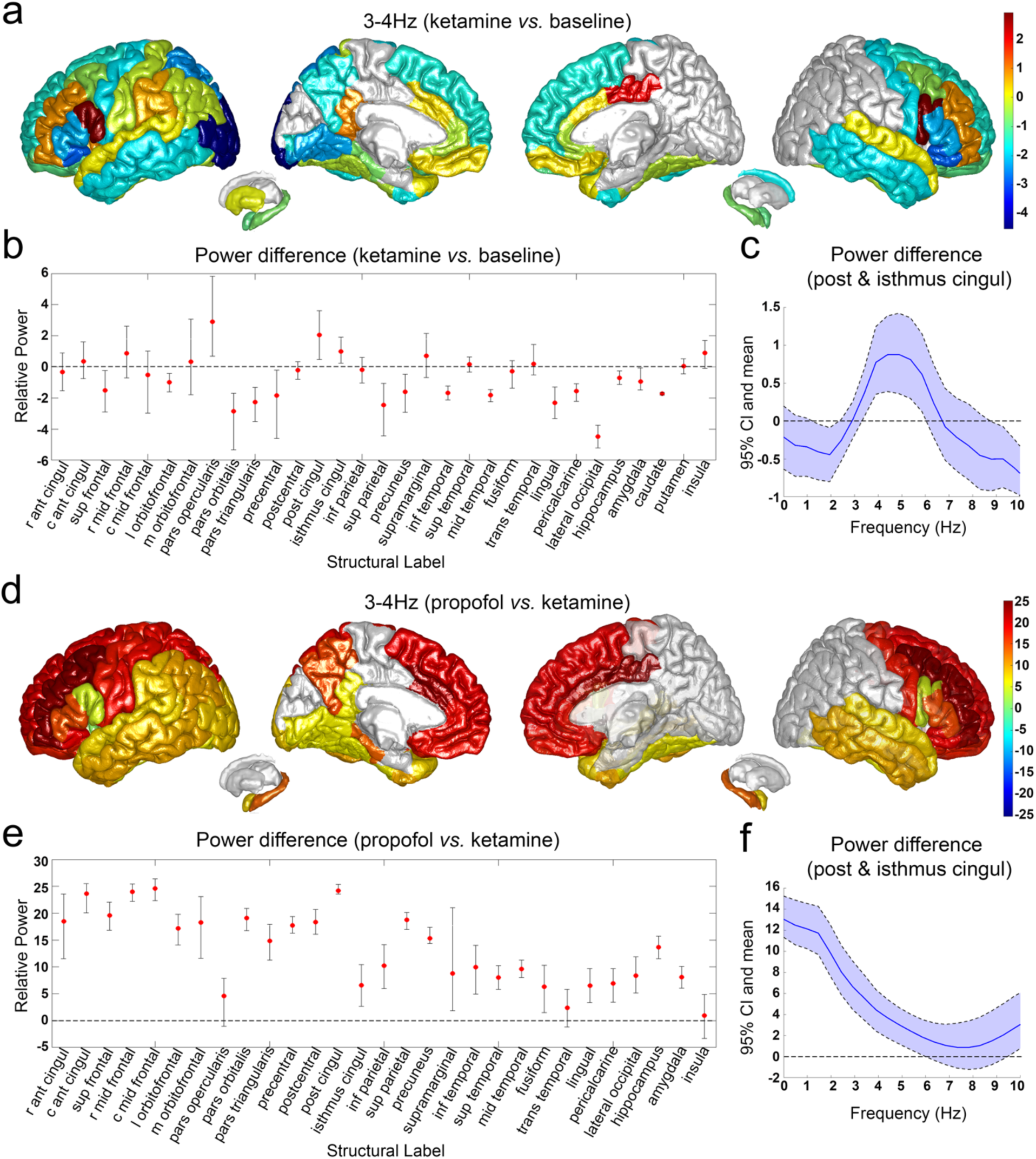
Analysis of 3-4Hz intracranial EEG power changes after administration of ketamine and propofol. **a**, structural mapping for 3-4Hz intracranial EEG power (dB) changes after ketamine infusion. **b**, mean and bootstrap 95% confidence interval (CI) for 3-4 Hz intracranial EEG power (dB) changes after ketamine infusion. **c**, power spectrum for posterior and isthmus cingulate cortex after ketamine infusion. **d**, structural mapping for 3-4Hz intracranial EEG power (dB) changes after propofol bolus. **e**, mean and bootstrap 95% confidence interval (CI) for 3-4 Hz intracranial EEG power (dB) changes after propofol bolus. **f**, power spectrum for posterior and isthmus cingulate cortex after propofol bolus. r ant cingul: rostral anterior cingulate; c ant cingul: caudal anterior cingulate; sup frontal: superior frontal; r mid frontal: rostral middle frontal; c mid frontal: caudal middle frontal; l orbitofrontal: lateral orbitofrontal; m orbitofrontal: medial orbitofrontal; post cingul: posterior cingulate; isthmus cingul: isthmus cingulate; inf parietal: inferior parietal; inf temporal: inferior temporal; sup temporal: superior temporal; mid temporal: middle temporal; trans temporal: transverse temporal.

**Figure 5.**
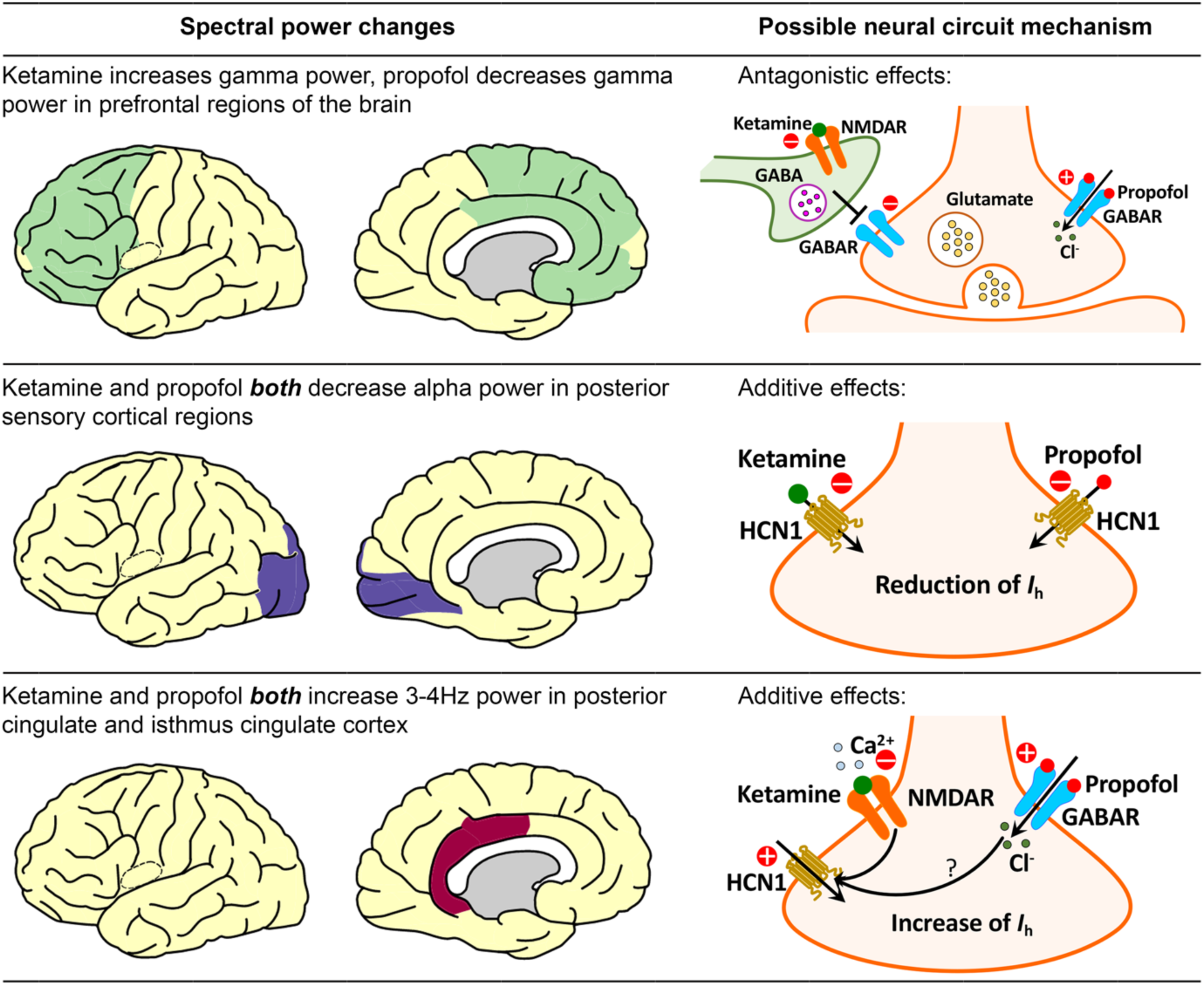
Possible neural circuit mechanisms for subanesthetic dose of ketamine and propofol-induced spectral power changes. NMDAR: N-methyl-D-aspartate receptor; GABA: gamma-aminobutyric acid; GABAR: gamma-aminobutyric acid receptor; HCN1: hyperpolarization-activated cyclic nucleotide-gated potassium channel 1; *Ih*: hyperpolarization-activated cationic current.

Besides its dissociative effects, subanesthetic ketamine has been shown to have a powerful anti-depressant effect. The oscillatory circuit dynamics produced by ketamine may be related to this anti-depressant effect. Subjects with a history of depression have been observed to have higher amplitude delta and theta oscillations compared to controls for negative targets in the backwards trial during a working memory task^34^. Consistent with this observation, we found that ketamine reduces delta and theta oscillation power (**Figure 2b&c**). Patients with depression have also been reported to have increased activity in alpha, beta, and theta bands at occipital and parietal regions of the brain^35^. Accordingly, we identified a global reduction of power at theta, alpha and beta frequencies, with the largest reduction in occipital and parietal regions after ketamine infusion (**Figure 2c,d&e**). Gamma oscillations have also been discussed as a potential biomarker for depression. Changes in gamma rhythms can vary according to behavioral states and task conditions, but there are a few studies suggesting that low gamma power is associated with depression. One EEG study found that subjects with high depression scores had reduced resting gamma in the anterior cingulate cortex^36^. Another MEG study showed that depressed subjects with lower baseline gamma and higher ketamine-induced gamma had a better response to ketamine than those with higher baseline gamma^37^. It has also known that the prefrontal cortex and hippocampus are implicated in ketamine’s antidepressant response^22^. The dramatic increase in gamma rhythms we identified in those brain regions with subanesthetic doses of ketamine are consistent with previous studies (**Figure 2f**).

Through this study, we characterized brain dynamics in cortical and subcortical structures during subanesthetic ketamine-induced dissociation using iEEG. We also analyzed drug interactions between ketamine and propofol to characterize the role of NMDA receptor disinhibition and HCN1 inhibition in different cortical and subcortical networks. We found that the subanesthetic ketamine-induced dissociative state was accompanied with a remarkable increase of gamma power in frontal regions of the brain and a large reduction of alpha power in occipital regions. The presence of propofol dramatically reversed the increase of frontal gamma power, implying an NMDA disinhibitory mechanism for ketamine that could be reversed by enhanced GABA inhibition. Meanwhile propofol further intensified the posterior reduction of alpha power, which could imply a shared HCN1 inhibitory mechanism. The gamma power increase we identified in prefrontal cortical areas and the hippocampus may underlie ketamine’s antidepressant effects. The alpha power decrease spanning posterior cortical regions is similar to propofol and may represent a common mechanism by which anesthetic drugs disrupt sensory contents of consciousness. In addition, we showed that a 3 Hz rhythm develops in the PMC during ketamine-induced dissociation in humans, consistent with previous studies in mice^23^. NMDA receptors, GABA receptors, and HCN1 channels may engage together to mediate the 3 Hz oscillation identified at PMC. The neural circuit mechanisms underlying ketamine-induced oscillatory dynamics, and their potential links to anti-depressive and dissociative effects as proposed in this study, may have important implications for the development of novel therapies with fewer side effects and greater safety. Future studies investigating brain dynamics after ketamine infusion in depressed patients are needed. Our results also show how the combination of ketamine and propofol could contribute to unconsciousness through a shared mechanism, providing an explanation for why propofol and ketamine appear to work synergistically to maintain unconsciousness when administered during general anesthesia^38^. Overall, we find that ketamine has distinct dynamic effects on neural systems known to mediate cognition, depression, and sensory processing mediated by multiple dissociable neuropharmacological mechanisms.

## Methodology

### Subject recruitment

Patients with medication-refractory epilepsy implanted with intracranial depth electrodes to locate their seizure onset zone were recruited from Massachusetts General Hospital and Brigham and Women’s Hospital. Electrode placement was determined by the clinical team independent of this study. Ten patients (five male and five female) aged 22 to 59 years old were recruited. Subjects’ demographic and electrode information are summarized in **Extended Data Table 1**. This study was approved by the Institutional Review Board (IRB) covering the two hospitals (Mass General Brigham Human Research Committee). Informed consent was obtained from all subjects prior to the study.

### Experimental procedure

All experiments were conducted during stereotactic neurosurgery for removal of the intracranial depth electrodes in the operating room at the Massachusetts General Hospital or the Brigham and Women’s Hospital. Participants were implanted with multi-lead depth electrodes (a.k.a. stereotactic EEG, sEEG) to confirm the hypothesized seizure focus, and located epileptogenic tissue in relation to essential cortex, thus directing surgical treatment. Depth electrodes (Ad-tech Medical, Racine WI, USA, or PMT, Chanhassen, MN, USA) with diameters of 0.8-1.0 mm and consisting of 8-16 platinum/iridium-contacts 1-2.4 mm long were stereotactically placed in locations deemed necessary for seizure localization by a multidisciplinary clinical team. The first period was a baseline recording of 5 minutes **(Figure 1a)**. The second period consisted of 14 minutes with continuous infusion of subanesthetic level of ketamine (total dose of 0.5mg/kg over 14 minutes). At the end of ketamine infusion, patients completed the Clinician-Administered Dissociative States Scale (CADSS) questionnaire^1^. Because of limited time in the operating room, patients were instructed to only answer yes or no to the questions. Immediately after the questionnaire, propofol bolus was given to the patients to induce general anesthesia. During the whole process, subjects were instructed to close their eyes to avoid eye-blink artifacts in the signal. Intracranial EEG (iEEG) signals were recorded using a Blackrock Cerebus system (Blackrock Microsystems) sampled at 2,000 Hz. Before each study, structural MRI scans were acquired for each subject (Siemens Trio 3 Tesla, T1-weighted magnetization-prepared rapid gradient echo, 1.3-mm slice thickness, 1.3×1 mm in-plane resolution, TR/TE=2530/3.3 ms, 7° flip angle).

### iEEG preprocessing and power spectral analysis

Raw iEEG data were notch filtered at 60Hz and its harmonics, downsampled to 500Hz, and detrended across the entire recording. The signals were then visually inspected, and channels with noise or artifacts were removed. Data were re-referenced with a bipolar montage. A total of 824 bipolar channels were generated for 10 subjects received ketamine, and 606 bipolar channels were generated for 7 subjects received propofol **(Extended Data Figure 1)**. Spectral analysis was performed using the multitaper method, with window lengths of T=2 sec with 0.5 sec overlap, time-bandwidth product TW=3, number of tapers K=5, and spectral resolution of 3 Hz^2,3^. The mean power spectral density for baseline, ketamine and propofol conditions were calculated by taking the average across each period. The power spectral density was converted to decibels (dB) to facilitate easier comparisons. The differences of power after ketamine infusion relative to baseline, and propofol relative to ketamine periods were calculated by subtracting the mean power spectral density in dB between each of the two conditions at different frequencies (slow: 0.1-1Hz, delta: 1-4Hz, theta: 4-8Hz, alpha: 8-15Hz, beta: 15-25Hz, gamma: 25-55Hz, low gamma: 25-40Hz, high gamma: 40-55Hz). Bootstrap method was used to generate the 95% confidence interval around the mean differences in power for each structural label at each frequency. The upper and lower bars represent the bootstrapped 95% confidence interval bounds. *P* < 0.05 was considered as statistically significant.

### Structural parcellation of the brain

The electrode positions in each subject’s brain were obtained by aligning the preoperative T1-weighted MRI with a postoperative CT/MRI using the Freesurfer image analysis tool^4,5^. To identify the structural label and functional network for each of the electrodes, an electrode labeling algorithm (ELA) was employed^6^. This algorithm estimated the probability of overlap of an expanding area around each electrode with brain structural labels that had been identified in the DKT40 atlas using purely anatomical approaches^7-11^. Then the ELA used gradient descent to find the closest voxel in the template’s brain that gives similar regions and probabilities to transform the patients’ electrode coordinates to the template brain^7-11^. Based on DKT40 atlas, we assigned the 824 electrodes from 10 subjects received ketamine to 49 structural labels, which were then further classified into 15 labels according to the anatomical locations and the mean differences of power after ketamine relative to baseline condition. Likewise, we assigned the 606 electrodes collected from 7 subjects received propofol to 14 structural labels. We plotted all electrodes on Colin 27 template brain with colors per parcellated brain region indicating the differences in power for ketamine infusion period relative to baseline, as well as for propofol bolus relative to ketamine infusion period for each of the frequencies.

## Data availability

The data that support the findings of this study are available on request from the corresponding author. The raw data are not publicly available due to restrictions relating to the per-participant imaging data currently containing information that could compromise the privacy of research participants.

## Code availability

The codes that support the findings of this study are available from the corresponding author upon request.

## Acknowledgments

This work was supported by NIH grants 1R01AG056015 (PLP), P01GM118269 (ENB) and the Tiny Blue Dot Foundation (PLP).

## Author contributions

ENB, SSC, and PLP conceived the idea of the project. LDL, DWZ, RAP, SSC, and PLP designed the experiments. FT, LDL, DWZ, GBV, ACP, RZ, LASCM, OJA, RAP, LSA, ENE, GRC, ZMW, RMR and SSC recruited the subjects and collected the data. FT, LDL, ACP, NP, and DS performed data analysis. FT and PLP wrote the manuscript. All authors approved the final version of the manuscript.

## Competing Interests

P.L.P. is an inventor on patents assigned to MGH related to brain monitoring, an inventor on a patent licensed to Masimo by Massachusetts General Hospital and a Co-founder of PASCALL Systems, Inc., a company developing closed-loop physiological control systems for anesthesiology.

E.N.B. is an inventor on patents assigned to MGH related to brain monitoring, an inventor on a patent licensed to Masimo by Massachusetts General Hospital and a Co-founder of PASCALL Systems, Inc., a company developing closed-loop physiological control systems for anesthesiology.

