## Supplemental Data for "Characterizing ketamine-induced dissociation using human intracranial neurophysiology: brain dynamics, network activity, and interactions with propofol"

**Extended Data Table 1.** Subject demographic and clinical information

| Subject | Gender | Age | Number of Electrodes | Number of Bipolar Channels | Baseline Duration (s) | Ketamine Duration (s) | Propofol Duration (s) |
|---|---|---|---|---|---|---|---|
| 1 | F | 59 | 10 | 55 | 366 | 840 | NA |
| 2# | M | 22 | 11 | 57 | 370 | 739 | 164 |
| 3# | F | 28 | 12 | 68 | 341 | 705 | 235 |
| 4 | M | 22 | 10 | 60 | 307 | 850 | NA |
| 5# | M | 48 | 14 | 77 | 201 | 840 | 57 |
| 6# | F | 34 | 9 | 65 | 300 | 840 | 402 |
| 7# | F | 22 | 10 | 104 | 300 | 840 | 50 |
| 8 | M | 48 | 9 | 102 | 300 | 840 | NA |
| 9# | M | 33 | 11 | 121 | 300 | 840 | 235 |
| 10# | F | 43 | 10 | 115 | 300 | 840 | 445 |

### Subjects received both ketamine and propofol; F: female, M: male; NA: not applicable

**Extended Data Table 2.** Summary of response to questionnaire

| Subject | Number of Yes (percentage) | Number of No (percentage) | Number of No Response (percentage) | Total |
|---|---|---|---|---|
| 1 | 8 (57.1%) | 4 (28.6%) | 2 (14.3%) | 14[#] |
| 2 | 16 (69.6%) | 7 (30.4%) | 0 (0%) | 23 |
| 3 | 16 (69.6%) | 7 (30.4%) | 0 (0%) | 23 |
| 4 | 5 (31.3%) | 0 (0%) | 11 (68.8%) | 16[#] |
| 5 | 7 (43.8%) | 2 (12.5%) | 7 (43.8%) | 16[#] |
| 6 | 5 (21.7%) | 18 (78.3%) | 0 (0%) | 23 |
| 7 | 16 (69.6%) | 7 (30.4%) | 0 (0%) | 23 |
| 8 | 21 (91.3%) | 0 (0%) | 2 (8.7%) | 23 |
| 9 | 16 (69.6%) | 7 (30.4%) | 0 (0%) | 23 |
| 10 | 2 (8.7%) | 21 (91.3%) | 0 (0%) | 23 |

[#] We did not finish all the questions for this patient because of limited time in the operating room

**Extended Data Table 3.** Mean and bootstrap 95% confidence interval for intracranial EEG power changes after ketamine infusion relative to baseline at 6 frequencies for 15 structural labels

| Structural Labels (N=10, n=824) | Slow (0.1-1Hz) | Delta (1-4Hz) | Theta (4-8Hz) | Alpha (8-15Hz) | Beta (15-25Hz) | Gamma (25-55Hz) |
|---|---|---|---|---|---|---|
| Anterior and Posterior Cingulate (N=7, n=55) | -2.06* [-2.96, -1.26] | -3.88* [-6.19, -1.71] | 2.41 [-0.05, 4.88] | -14.33* [-19.52, -9.11] | 4.49 [-2.58, 12.15] | 159.04* [134.42, 189.31] |
| Superior Frontal (N=7, n=23) | -2.27* [-3.87, -0.82] | -6.67* [-10.74, -2.83] | -5.65* [-11.94, -1.15] | -24.11* [-37.22, -11.36] | 0.54 [-13.91, 16.47] | 153.03* [115.95, 204.18] |
| Middle Frontal (N=7, n=79) | -2.80* [-3.97, -1.86] | -4.93* [-7.97, -2.30] | 0.02 [-4.18, 3.88] | -26.48* [-33.50, -20.14] | -14.26* [-21.34, -6.56] | 153.59* [130.80, 181.64] |
| Orbitofrontal (N=7, n=49) | 0.25 [-0.87, 1.65] | -2.83 [-5.27, 0.23] | 0.63 [-1.34, 2.58] | -7.14* [-11.21, -3.14] | 2.17 [-2.56, 7.72] | 133.68* [113.17, 162.77] |
| Inferior Frontal (N=9, n=56) | -2.54* [-4.36, -1.25] | -7.92* [-11.72, -5.04] | -0.44 [-3.39, 2.47] | -11.83* [-16.42, -8.10] | 4.63 [-0.01, 10.28] | 149.20* [126.44, 175.51] |
| Precentral (N=5, n=18) | -2.44* [-5.67, -0.87] | -8.09* [-15.67, -3.74] | -9.96* [-23.05, -2.58] | -26.50* [-53.55, -12.97] | -18.21* [-24.41, -12.16] | 91.70* [64.17, 138.70] |
| Postcentral (N=3, n=23) | -1.74* [-3.23, -0.50] | -4.68* [-7.45, -2.18] | -4.50* [-7.76, -1.03] | -33.55* [-40.24, -25.96] | -36.00* [-44.52, -26.68] | 72.47* [48.84, 96.83] |
| Isthmus Cingulate (N=4, n=25) | -0.61 [-1.50, 0.22] | -0.08 [-1.77, 1.36] | 1.44 [-2.08, 5.94] | -13.90* [-17.34, -9.92] | -10.53* [-14.76, -6.99] | 43.09* [34.43, 53.46] |
| Parietal, Precuneus and Supramarginal (N=4, n=45) | -2.40* [-2.99, -1.93] | -4.84* [-6.61, -3.55] | -7.75* [-10.71, -5.66] | -14.94* [-18.37, -12.23] | -15.90* [-20.59, -11.49] | 5.15 [-9.49, 15.96] |
| Temporal and Fusiform (N=10, n=260) | -2.34* [-2.71, -2.02] | -5.37* [-6.23, -4.63] | -5.83* [-7.39, -4.41] | -12.02* [-13.36, -10.76] | -8.40* [-10.37, -6.17] | 28.87* [22.35, 36.81] |
| Lingual and Pericalcarine (N=4, n=39) | -1.64* [-2.49, -0.77] | -6.84* [-8.63, -5.18] | -11.68* [-15.10, -7.64] | -27.02* [-31.36, -23.23] | -19.58* [-24.61, -13.98] | 19.07* [9.21, 30.70] |
| Occipital (N=2, n=18) | -3.51* [-4.29, -2.30] | -12.91* [-15.15, -10.13] | -20.52* [-23.12, -17.76] | -32.07* [-38.12, -24.93] | -43.81* [-48.21, -38.54] | -42.96* [-68.30, -18.75] |
| Hippocampus and Amygdala (N=9, n=113) | -1.51* [-1.96, -0.95] | -3.91* [-5.02, -2.36] | -0.25 [-2.09, 1.54] | -1.54 [-3.81, 0.83] | 4.43* [1.69, 7.64] | 68.64* [56.44, 82.26] |
| Striatum (N=3, n=4) | 0.62 [-2.51, 3.01] | -1.40* [-2.63, -0.16] | -1.85* [-2.94, -0.12] | -26.60* [-32.82, -17.48] | -2.47 [-10.32, 10.23] | 96.37* [53.39, 119.42] |
| Insula (N=4, n=17) | -0.23 [-1.17, 0.49] | 0.13 [-2.49, 2.02] | 3.88* [1.42, 6.56] | -5.44* [-8.42, -2.47] | 1.46 [-2.57, 7.51] | 56.30* [41.29, 75.39] |

N: number of subjects; n: number of electrodes; *: significant difference, $P < 0.05$

**Extended Data Table 4.** Mean and bootstrap 95% confidence interval for intracranial EEG power changes after propofol bolus relative to ketamine period at 7 frequencies for 14 structural labels

| Structural Labels (N=7, n=606) | Slow (0.1-1Hz) | Delta (1-4Hz) | Theta (4-8Hz) | Alpha (8-15Hz) | Beta (15-25Hz) | Low Gamma (25-40Hz) | High Gamma (40-55Hz) |
|---|---|---|---|---|---|---|---|
| Anterior and Posterior Cingulate (N=4, n=30) | 38.69* [33.40, 41.91] | 87.25* [74.65, 95.27] | 60.54* [50.71, 68.26] | 144.08* [128.50, 158.41] | 145.45* [116.86, 167.21] | -16.45 [-71.27, 29.72] | -61.26* [-99.98, -22.67] |
| Superior Frontal (N=4, n=14) | 33.66* [29.40, 37.45] | 77.24* [67.49, 86.04] | 52.74* [46.16, 59.77] | 124.91* [99.21, 152.48] | 108.56* [68.59, 162.20] | -32.50 [-119.83, 44.73] | -68.32* [-146.89, -17.11] |
| Middle Frontal (N=4, n=43) | 38.66* [35.20, 41.59] | 92.64* [86.29, 97.80] | 57.42* [51.52, 62.36] | 113.78* [99.02, 128.28] | 114.66* [91.13, 136.67] | -68.95* [-118.38, -21.95] | -134.51* [-177.93, -96.62] |
| Orbitofrontal (N=6, n=43) | 30.58* [26.17, 34.75] | 69.38* [59.03, 78.04] | 51.24* [44.36, 57.60] | 137.43* [121.60, 150.88] | 166.62* [139.86, 190.57] | 44.88 [-0.68, 82.50] | -52.27* [-93.71, -20.62] |
| Inferior Frontal (N=6, n=39) | 25.16* [20.11, 29.67] | 58.60* [46.56, 68.51] | 39.23* [31.56, 46.41] | 110.97* [91.94, 127.35] | 155.03* [124.18, 181.32] | 27.40 [-16.10, 63.85] | -61.86* [-107.45, -24.86] |
| Precentral (N=4, n=17) | 25.03* [20.92, 30.05] | 65.87* [58.90, 74.49] | 63.91* [59.31, 70.28] | 158.32* [141.96, 179.07] | 175.00* [136.46, 212.71] | 136.59* [80.92, 197.51] | 49.39* [12.05, 91.84] |
| Postcentral (N=2, n=5) | 18.30* [14.43, 23.12] | 57.76* [49.72, 66.49] | 68.88* [64.37, 78.00] | 154.79* [127.73, 179.99] | 214.90* [186.07, 241.30] | 170.25* [135.60, 223.74] | 67.34* [32.12, 97.19] |
| Isthmus Cingulate (N=3, n=20) | 20.69* [17.61, 24.98] | 33.62* [23.21, 45.70] | 8.12 [-6.46, 21.40] | 33.33* [14.49, 54.32] | 98.33* [79.28, 117.16] | 86.48* [73.32, 99.47] | 9.78* [0.02, 17.19] |
| Parietal, Precuneus and Supramarginal (N=3, n=35) | 26.19* [23.53, 28.48] | 55.35* [46.56, 62.83] | 22.58* [14.76, 30.52] | 34.81* [23.51, 49.37] | 82.40* [72.07, 95.51] | 56.20* [47.10, 64.57] | 4.06 [-2.04, 10.02] |
| Temporal and Fusiform (N=7, n=206) | 24.00* [22.42, 25.64] | 42.53* [38.55, 46.45] | 23.13* [19.15, 26.86] | 45.74* [39.17, 52.19] | 97.43* [88.09, 106.81] | 64.81* [53.53, 75.52] | 1.33 [-9.12, 10.38] |
| Lingual and Pericalcarine (N=3, n=33) | 19.33* [16.72, 22.27] | 33.42* [25.40, 41.48] | 1.87 [-8.90, 12.02] | -15.66 [-38.74, 4.61] | 12.61 [-5.12, 29.70] | 44.45* [33.00, 55.02] | -4.32 [-15.44, 6.87] |
| Occipital (N=2, n=18) | 24.75* [21.21, 29.22] | 43.60* [33.34, 55.91] | -3.32 [-13.56, 8.03] | -35.20* [-58.21, -14.02] | 22.00* [1.95, 42.76] | 60.47* [50.09, 71.30] | 18.85* [11.40, 26.00] |
| Hippocampus and Amygdala (N=7, n=90) | 20.87* [18.74, 23.05] | 43.87* [39.02, 48.67] | 43.99* [38.04, 49.77] | 111.87* [100.95, 122.29] | 151.28* [137.11, 164.07] | 108.87* [91.29, 124.79] | 20.56* [4.73, 34.94] |
| Insula (N=3, n=13) | 7.31* [4.67, 11.50] | 10.29* [0.37, 21.78] | 17.35* [4.97, 30.18] | 116.90* [99.23, 142.82] | 124.26* [105.76, 149.45] | 73.51* [52.86, 98.92] | 8.17 [-0.07, 20.66] |

N: number of subjects; n: number of electrodes; *: significant difference, $P < 0.05$

**Extended Data Table 5.** Mean and bootstrap 95% confidence interval for 3-4 Hz intracranial EEG power changes after administration of ketamine and propofol

| Structural Labels | Ketamine vs. Baseline (N=10, n=824) | Propofol vs. Ketamine (N=7, n=606) |
|---|---|---|
| Rostral Anterior Cingulate | -0.33 [-1.52, 0.88] (N=7, n=26) | 18.48 [11.64, 23.45]* (N=4, n=13) |
| Caudal Anterior Cingulate | 0.35 [-0.81, 1.57] (N=6, n=25) | 23.62 [20.37, 25.57]* (N=3, n=13) |
| Superior Frontal | -1.51 [-2.87, -0.25]* (N=7, n=23) | 19.56 [16.92, 22.11]* (N=4, n=14) |
| Rostral Middle Frontal | 0.86 [-0.70, 2.62] (N=7, n=57) | 24.01 [22.29, 25.43]* (N=4, n=32) |
| Caudal Middle Frontal | -0.53 [-2.88, 1.08] (N=6, n=22) | 24.57 [22.26, 26.37]* (N=3, n=11) |
| Lateral Orbitofrontal | -0.98 [-1.61, -0.41]* (N=7, n=39) | 17.18 [14.11, 19.81]* (N=6, n=33) |
| Medial Orbitofrontal | 0.34 [-1.89, 2.99] (N=6, n=10) | 18.27 [11.78, 23.27]* (N=6, n=10) |
| Pars Opercularis | 2.90 [0.60, 5.82]* (N=4, n=10) | 4.58 [-1.02, 7.92] (N=2, n=3) |
| Pars Orbitalis | -2.86 [-5.30, -1.68]* (N=5, n=7) | 19.07 [16.75, 20.93]* (N=4, n=6) |
| Pars Triangularis | -2.25 [-3.54, -1.33]* (N=7, n=39) | 14.86 [11.19, 17.85]* (N=5, n=30) |
| Precentral | -1.84 [-4.61, -0.21]* (N=5, n=18) | 17.70 [16.38, 19.46]* (N=4, n=17) |
| Postcentral | -0.22 [-0.81, 0.32] (N=3, n=23) | 18.37 [16.05, 20.68]* (N=2, n=5) |
| Posterior Cingulate | 2.05 [0.48, 3.61]* (N=1, n=4) | 24.18 [23.47, 25.36]* (N=1, n=4) |
| Isthmus Cingulate | 1.00 [0.24, 1.90]* (N=4, n=25) | 6.58 [2.75, 10.29]* (N=3, n=20) |
| Inferior Parietal | -0.19 [-1.01, 0.60] (N=3, n=24) | 10.22 [5.90, 14.11]* (N=2, n=18) |
| Superior Parietal | -2.45 [-4.41, -1.03]* (N=1, n=9) | 18.72 [16.92, 20.16]* (N=1, n=9) |
| Precuneus | -1.60 [-2.92, -0.48]* (N=1, n=5) | 15.30 [14.40, 17.37]* (N=1, n=5) |
| Supramarginal | 0.71 [-0.72, 2.10] (N=3, n=7) | 8.80 [1.83, 21.03]* (N=2, n=3) |
| Inferior Temporal | -1.68 [-2.14, -1.23]* (N=6, n=29) | 9.95 [4.82, 13.90]* (N=5, n=24) |
| Superior Temporal | 0.16 [-0.32, 0.64] (N=10, n=74) | 8.04 [5.86, 10.21]* (N=7, n=51) |
| Middle Temporal | -1.82 [-2.23, -1.45]* (N=10, n=141) | 9.65 [8.01, 11.26]* (N=7, n=115) |
| Fusiform | -0.28 [-1.34, 0.43] (N=4, n=10) | 6.34 [1.64, 10.35]* (N=4, n=10) |
| Transverse Temporal | 0.19 [-0.53, 1.49] (N=2, n=6) | 2.39 [-1.20, 5.81] (N=2, n=6) |
| Lingual | -2.31 [-3.29, -1.28]* (N=4, n=33) | 6.52 [3.35, 9.71]* (N=3, n=27) |
| Pericalcarine | -1.56 [-2.25, -1.10]* (N=1, n=6) | 6.93 [3.48, 9.69]* (N=1, n=6) |
| Lateral Occipital | -4.49 [-5.21, -3.76]*(N=2, n=18) | 8.36 [5.10, 11.85]* (N=2, n=18) |
| Hippocampal | -0.70 [-1.13, -0.24]* (N=9, n=72) | 13.70 [11.53, 15.75]* (N=7, n=58) |
| Amygdala | -0.95 [-1.49, -0.03]* (N=8, n=41) | 8.08 [6.06, 10.19]* (N=6, n=32) |
| Caudate | -1.73 [-1.81, -1.65]* (N=1, n=2) | NA |
| Putamen | 0.04 [-0.46, 0.53] (N=2, n=2) | NA |
| Insula | 0.89 [-0.11, 1.66] (N=4, n=17) | 0.95 [-3.11, 4.89] (N=3, n=13) |

N: number of subjects; n: number of electrodes; NA: not applicable; *: significant difference, $P < 0.05$

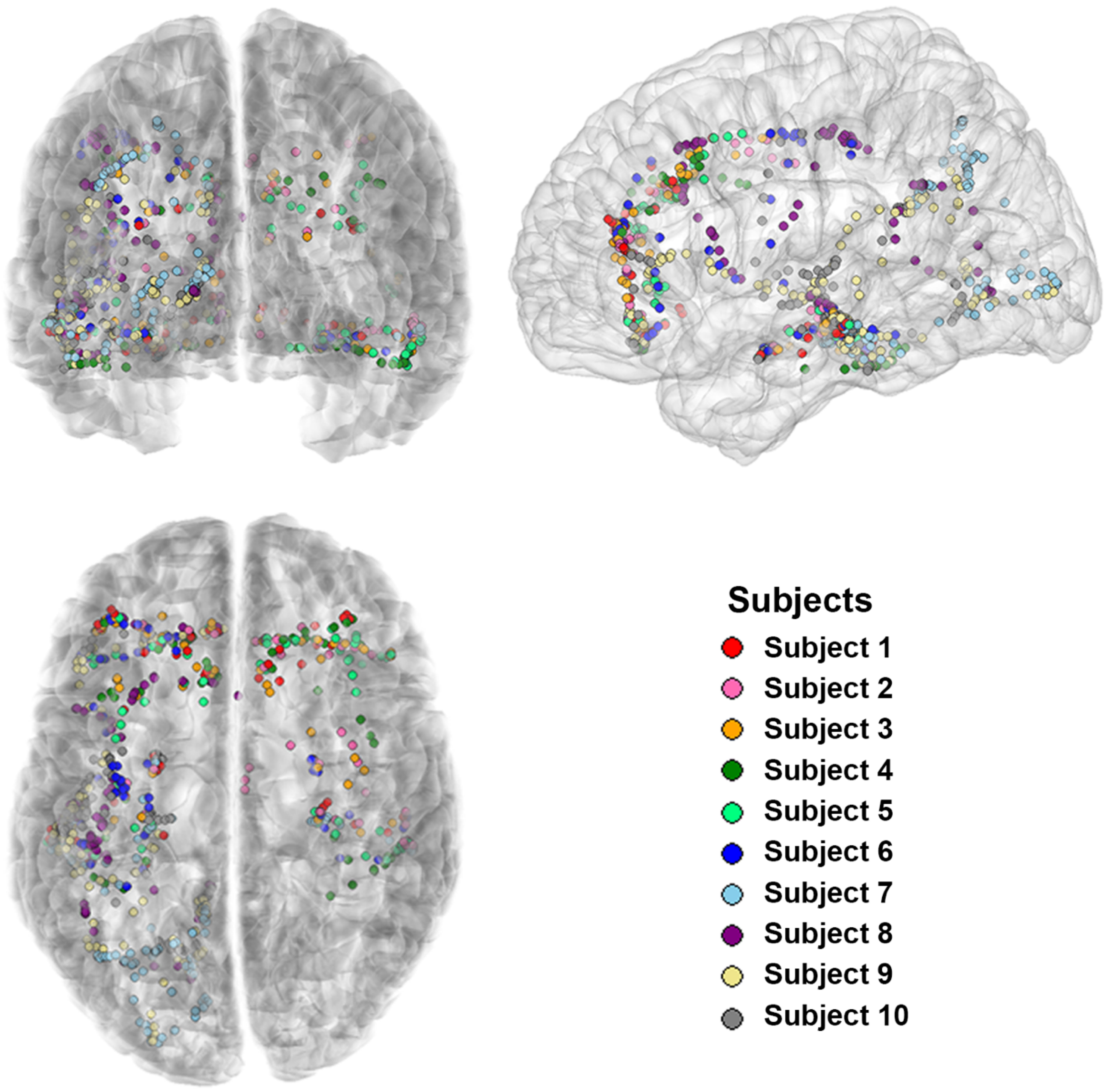


**Extended Data Figure 1.** Coregistered and reconstructed intracranial electrodes from all subjects on Colin 27 brain template

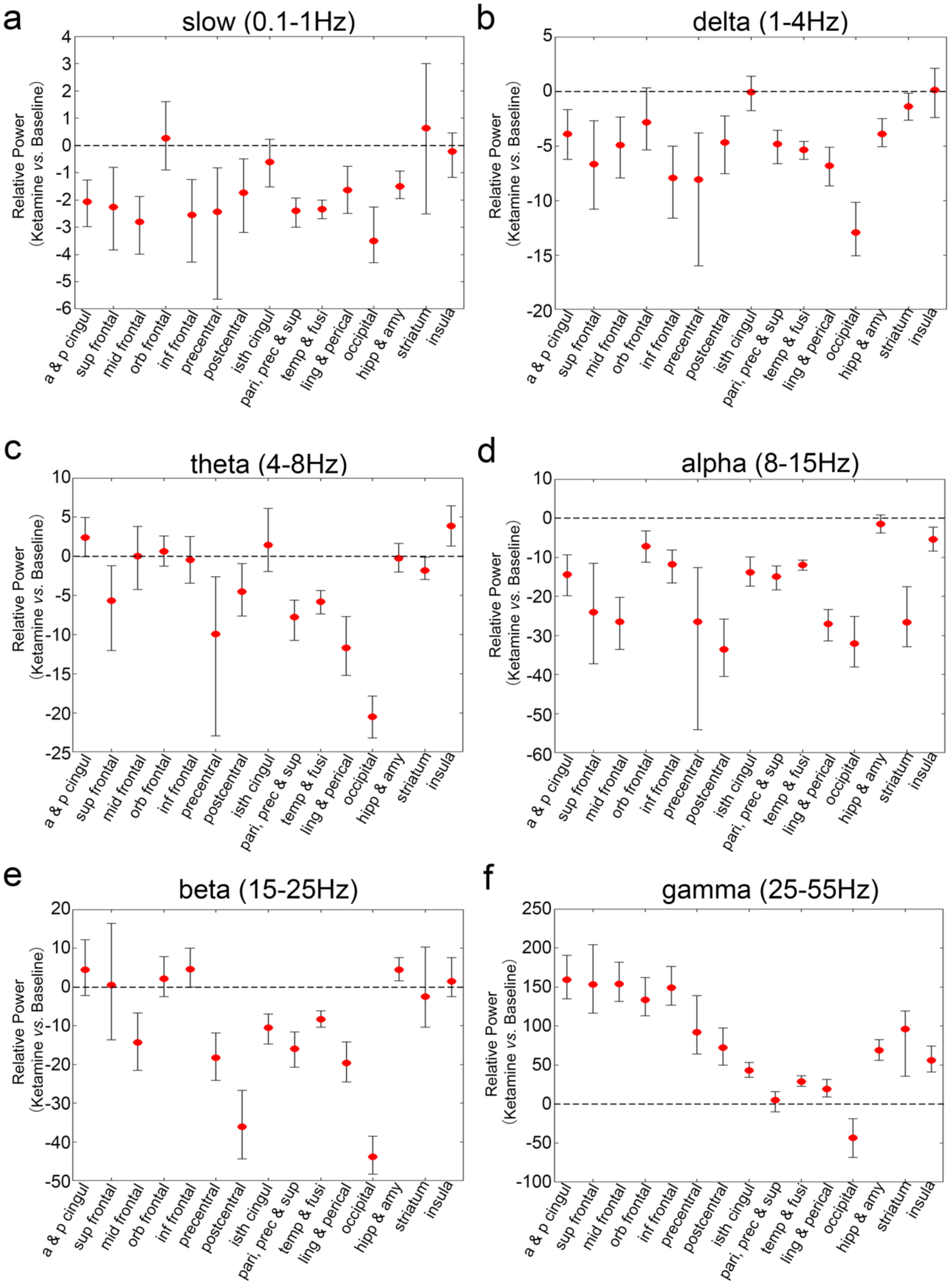


**Extended Data Figure 2.** Mean and bootstrap 95% confidence interval for intracranial EEG power changes after ketamine infusion relative to baseline at 6 frequencies for 15 structural labels: a & p cingul (anterior and posterior cingulate), sup frontal (superior frontal), mid frontal (middle frontal), orb frontal (orbitofrontal), inf frontal (parsopercularis, parsorbitalis, and parstriangularis), precentral, postcentral, isth cingul (isthmuscingulate), pari, prec & sup (parietal, precuneus, and supramarginal), temp & fusi (temporal and fusiform), ling & perical (lingual and pericalcarine), occipital, hipp & amy (hippocampal and amygdala), striatum (caudate and putamen), and insula.

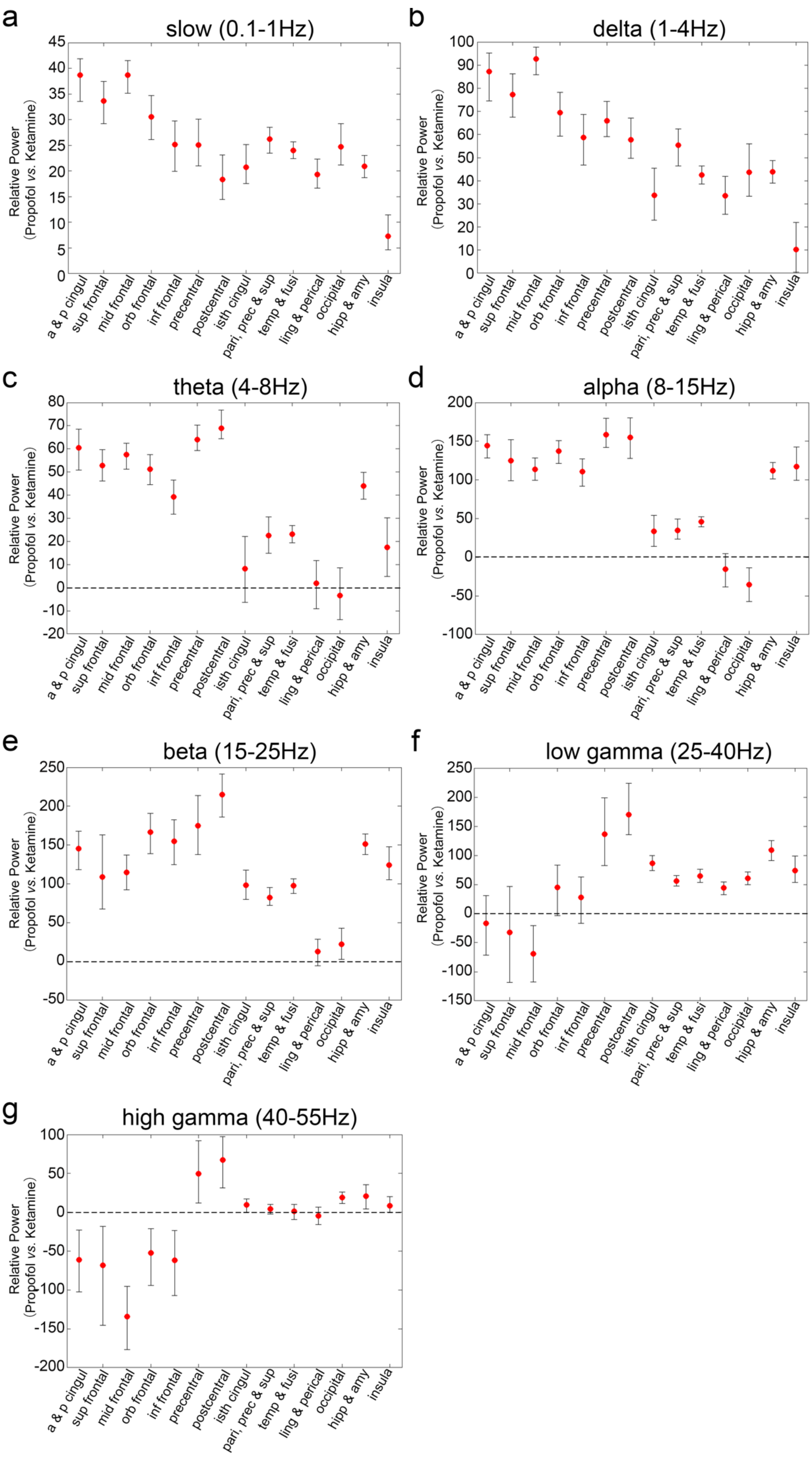
a
slow (0.1-1Hz)
b
delta (1-4Hz)
c
theta (4-8Hz)
d
alpha (8-15Hz)
e
beta (15-25Hz)
f
low gamma (25-40Hz)
g
high gamma (40-55Hz)
Relative Power
(Propofol vs. Ketamine)
a & p cingul
sup frontal
mid frontal
orb frontal
inf frontal
precentral
postcentral
isth cingul
pari, prec & sup
temp & fusi
ling & perical
occipital
hipp & amy
insula

**Extended Data Figure 3.** Mean and bootstrap 95% confidence interval for intracranial EEG power changes after propofol bolus relative to ketamine period at 7 frequencies for 14 structural labels: a & p cingul (anterior and posterior cingulate), sup frontal (superior frontal), mid frontal (middle frontal), orb frontal (orbitofrontal), inf frontal (parsopercularis, parsorbitalis, and parstriangularis), precentral, postcentral, isth cingul (isthmuscingulate), pari, prec & sup (parietal, precuneus, and supramarginal), temp & fusi (temporal and fusiform), ling & perical (lingual and pericalcarine), occipital, hipp & amy (hippocampal and amygdala), and insula.
